# Expression of the transcription factor Isl1 in dopaminergic neurons of the mouse forebrain

**DOI:** 10.1101/2023.12.27.573451

**Authors:** Adriana C. Camarano, Marcelo Rubinstein, Flavio S. J. de Souza

## Abstract

The development of the bewildering assortment of neuronal types found in the vertebrate central nervous system (CNS) depends on the distribution of transcription factors and signalling molecules along the embryonic neural tube. The *Islet-1* (*Isl1*) gene, which encodes a transcription factor of the LIM-homeodomain family, is known to be expressed in the nervous system, playing crucial funtions in the differentiation of neuronal populations located in the spinal cord, striatum, hypothalamus and retina. Here, we use immunofluorencence to trace the distribution of Isl1 protein during the development of the mouse forebrain, with an emphasis on the hypothalamic area and its neighbouring regions. Isl1 is abundantly expressed in the subpallium, most of the hypothalamus and in the prethalamus. Interestingly, we found that Isl1 is expressed in most dopaminergic neurons of the forebrain in early development (e10.5, e11.5), as revealed by colabelling with the enzyme tyrosine hydroxylase (TH). At later stages (e18.5) and adulthood, the degree of colocalisation of Isl1 with TH decreases, but the factor is still found in most dopaminergic neurones of the forebrain, in particular of the prethalamic region (A13 group), tuberal hypothalamus (A12), preoptic area (A15) and part of the periventricular area (part of the A14 group). Altogether, our observations indicate that Isl1 is a molecular marker of forebrain dopaminergic groups and might play a role in the development of these neuronal populations.

## Introduction

Transcription factors are key regulators of gene activity, and mapping the expression of these proteins during embryogenesis is crucial to the understanding of how form and function are established during development. The transcription factor Islet-1 (Isl1) belongs to the LIM-HD family, characterised by the presence of a LIM domain, which mediates protein-protein interactions, and a DNA-binding homeodomain (HD), in addition to other functional domains (Karlsson et al, 1990; Rétaux and Bachy, 2002; Hunter and Rhoades, 2005; Bhati et al, 2008). Like other LIM-HD proteins, Isl1 is an ancient factor, found in all metazoan clades (Srivastava et al, 2010). In the mouse, the expression of Isl1 has been extensively studied in several regions of the central and peripheral nervous system, the retina, the developing heart, pancreas, thyroid and pituitary glands, urogenital area and limbs, among other regions (Thor et al, 1991; Ericson et al, 1992; Cai et al, 2003; Elshatory et al, 2007a; Zhuang et al, 2013; Bejarano-Escobar et al, 2015; Zhang et al, 2017). Studies with mice carrying mutant alleles of *Isl1* have revealed various functions for the factor during organogenesis. In the nervous system, *Isl1* is necessary for motorneuron differentiation in the spinal cord (Pfaff et al, 1996; Lee and Pfaff, 2003; Thaler et al, 2004), and is also necessary for proper differentiation of neuronal types in sensory ganglia (Sun et al, 2008), retina (Elshatory et al, 2007b; Elshatory et al, 2008; Bejarano-Escobar et al, 2015) and subpallium (Elshatory and Gan, 2008; Ehrman et al, 2013 Lu et al, 2014). In the tuberal hypothalamus, *Isl1* is necessary for the differentiation of neurons that produce the neuropeptides Proopiomelanocortin (Pomc; Nasif et al, 2015), AgRP, Somatostatin (Sst) and Growth hormone-releasing hormone (Grhr) (Lee et al, 2016).

Populations of dopaminergic neurons distributed in the midbrain (classified as groups A8-A10) and forebrain (groups A11-A16) can be identified by the expression of tyrosine hydroxylase (TH), the limiting enzyme necessary for dopamine synthesis (Marín et al, 2005, Björklund and Dunnett, 2007; Bilbao et al, 2022). In the forebrain, dopaminergic groups are located in the diencephalon (groups A11 and A13), hypothalamus (groups A12 and A14) preoptic area of the basal telencephalon (A15) and olfactory bulb (A16), although this traditional classification in groups does not account for the spread out distribution of TH+ neurons, which do not respect rigid boundaries between brain areas (see for instance Bilbao et al, 2022). For some authors, the A13 groups lies within the caudal part of the peduncular hypothalamus (Puelles et al, 2012; Bilbao et al, 2022). Double immunohistochemistry has revealed colocalisation of Isl1 and TH in some dopaminergic pupulations. Thus, Isl1 is found during development in the A13 group, located in the Zona Incerta (ZI) of the prethalamic region (Mastick and Andrews, 2001; Andrews et al, 2003; Espana and Clotman, 2012). Colocalisation of Isl1 and TH has also been observed in the arcuate nucleus (ARC) of the hypothalamus, corresponding to the A12 group (Lee et al, 2016).

In the forebrain, the expression of Isl1 has been extensively characterised in model vertebrates like the frog, *Xenopus laevis* (Moreno et al, 2008; Domínguez et al, 2013, 2014, 2015) and the zebrafish, *Danio rerio* (Schredelseker and Driever, 2020; López et al, 2021; Lozano et al, 2023), but a systematic description of Isl1 expression in the mouse forebrain is lacking. Here, we study the distribution of Isl1 in the hypothalamus and neighbouring areas (preoptic area and prethalamus) in the mouse and find a high degree of colocalisation of Isl1 protein in dopaminergic neuronal populations during development and adulthood. We employ the prosomeric model for forebrain development, which better reflects the general anatomical organisation of the brain and patterns of gene expression (Puelles and Ferrán, 2012; Puelles et al, 2013; Puelles and Rubenstein, 2015).

## Materials and Methods

### Animal husbandry

Mice of mixed genetic background were housed in ventilated cages under controlled temperature and photoperiod (12-h light/12-h dark cycle). All mouse procedures followed the Guide for the Care and Use of Laboratory Animals (Committee on Care and Use of Laboratory Animals,1996) and were in agreement with the institutional animal care and use committee at the Faculty of Exact and Natural Sciences, University of Buenos Aires.

### Embryo and brain processing and histology

Embryos were obtained and processed as in Orquera et al (2016). Briefly, pregnant females were sacrificed at the indicated stages and embryos were collected, washed in cold phosphate-buffered saline (PBS) and fixed for 30 min to 2h in 4% paraformaldehyde (PFA) in PBS at 4°C. After washing, samples were cryoprotected with 10% sucrose-PBS overnight. Tissues were then stabilised in 10% sucrose-10% gelatin-PBS at 37°C for 30 min prior to freezing. Gelatin blocks containing the tissues were placed on cork sheets with OCT compound (Biopack). Gelatin blocks were snap-frozen in 2-Methylbutane (Sigma, M32631) at −60°C and stored at −80°C. Tissue slices of 16–20 mm were cut in a cryostat (Leica CM1850). Anaesthesised adult mice were processed by cardiac perfusion and their brains processed was described (Nasif et al, 2015).

### Immunofluorescence and antibodies

Immunofluorescence was performed as described (Nasif et al, 2015). Primary antibodies used were: mouse anti-Isl1/2 (1:200; 40.2D6, Developmental Studies Hybridoma Bank, University of Iowa); rabbit monoclonal anti-Isl1 (1:1000; ab109517, Abcam); rabbit polyclonal anti-Isl1 (1:1000; ab10670, Abcam); goat anti-Isl1 (1:1000; R&D Systems AF1837); mouse anti-Nkx2.2 (1:1000, 74.5A5, Developmental Studies Hybridoma Bank, University of Iowa); rabbit anti-Nkx2.1 (1:1000, ab40880, Abcam); goat anti-Brn2 (1:250; C-20, Santa Cruz Biotechnology); goat anti-Lhx1 (1:500, sc-19341, Santa Cruz Biotechnology); rabbit anti-Pax6 (1:500; Poly19013, Biolegends); rabbit anti-TH (1:1000; 657012, Calbiotech); chicken anti-TH (1:1000; ab76442). Secondary antibodies were (all from Molecular Probes and used 1:1000): donkey anti-rabbit IgG AlexaFluor 555; donkey anti-goat IgG AlexaFluor 555; donkey anti-chicken IgG AlexaFluor 488; goat anti-rabbit IgG AlexaFluor 488).

### Fluorescence microscopy and quantifications

For epifluorescence microscopy an Olympus BX51 was used; confocal microscopy images were obtained with Olympus FluoView FV300, FV1000 and a Zeiss LSM900. Confocal images of sections spanning each anatomical region were analysed to estimate TH/Isl1 colocalisation. The number of TH+ neurons with Isl1+ nuclei and the total number of TH+ neurons from each region were counted for at least three embryos/brains for each stage. Means are presented with the standard deviation (SD).

## Results

### Expression of Isl1 in the developing forebrain

To map the expression of Isl1 during mouse development, we performed single and double immunofluorescence experiments with antibodies against Isl1 and other relevant molecular markers. In interpreting our results, we employ the prosomeric model of forebrain development, which better reflects the general anatomical organisation of the brain and patterns of gene expression (Puelles and Ferrán, 2012; Puelles et al, 2013; Ferran et al, 2015; Puelles and Rubenstein, 2015). We concentrated our analysis on the hypothalamus and surrounding areas, where several dopamine neuronal groups are known to be located.

At e10.5, near the start of neurogenesis, Isl1 expression is already abundant in the forebrain. At this stage, the expression domain of the homedodomain (HD) transcription factor Nkx2.2 labels the neuroepithelium of the hypothalamus and diencephalon along the alar/basal boundary. Within the diencephalon, Nkx2.2 delineates the Zona Limitans Intrathalamica (ZLI), which demarcates the limit between prosomeres 2 and 3 (Shimamura et al, 1995). Double immunofluorescence of Isl1/Nkx2.2 shows that Isl1 is mostly expressed in the basal plate of the hypothalamus, near the alar/basal limit labelled by Nkx2.2 (Fig. 1). In the alar plate, Isl1 is expressed in most of the mantle zone of diencephalic prosomere 3, which corresponds to the prethalamus (Pth). Only a few Isl1+ cells are observed posterior to the ZLI, within prosomere 2 (arrow in Fig. 1B). In the telencephalon, Isl1 is coexpressed with Nkx2.2 in the mantle zone of the preoptic area (POA) and the medium ganglionic eminence (MGE) in the subpallium (SPall). At E10.5, the HD transcription factor Nkx2.1 is expressed in most of the basal plate of the hypothalamus, as well as the POA, and most Isl1+ cells are located within the Nkx2.1-expression domain (Fig. 2). An exception is the prethalamus, where Nkx2.1 is not expressed (Fig. 2A).

**Fig 1:**
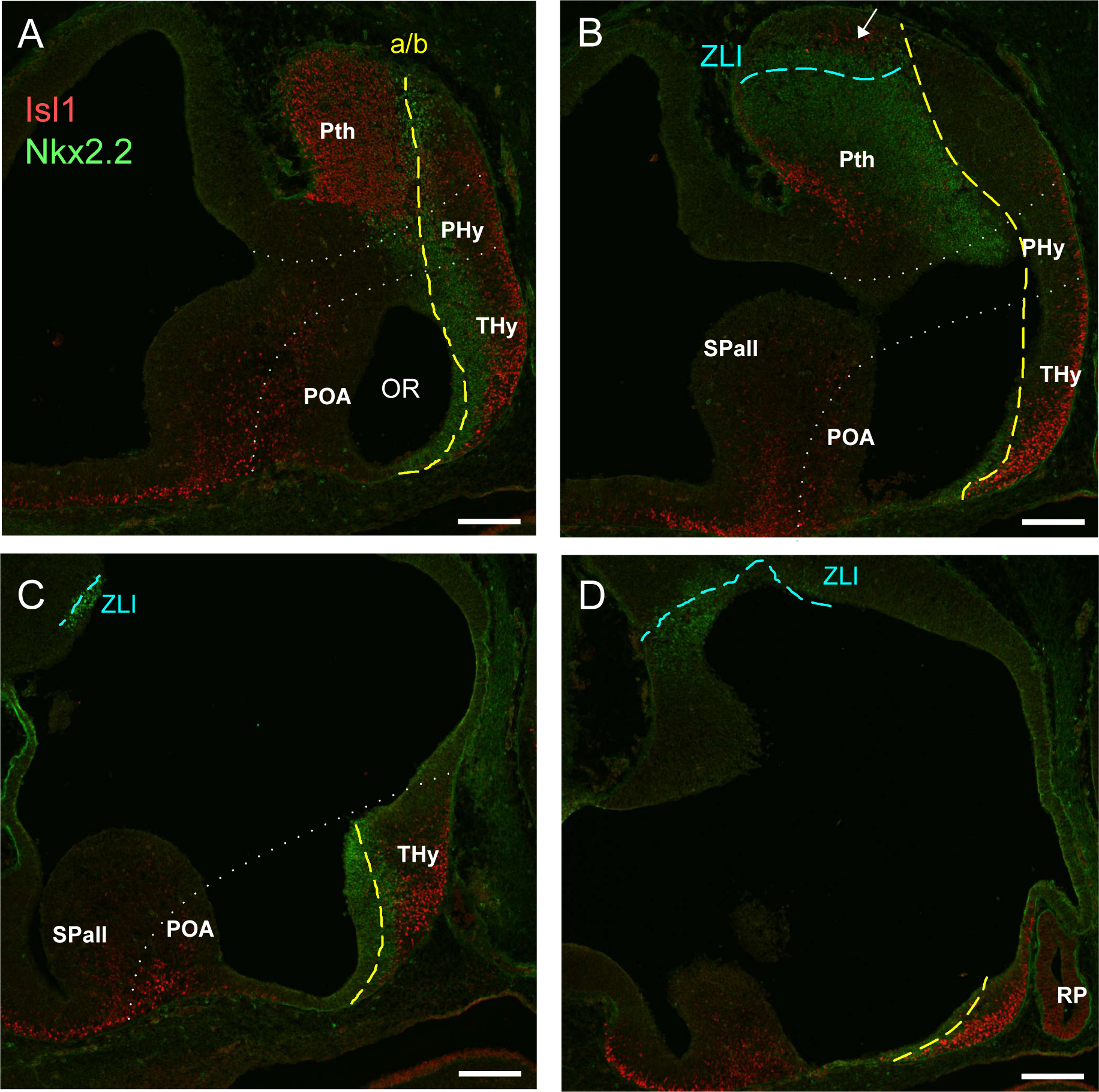
Double immunofluorescence of Isl1 (red) and Nkx2.2 (green) in sagital slices of a E10.5 mouse embryo. Panels go from lateral (A) to medial (D). Dotted white lines indicate the limits of anatomical regions. The yellow dotted line indicates the alar/basal limit (a/b).

**Fig 2:**
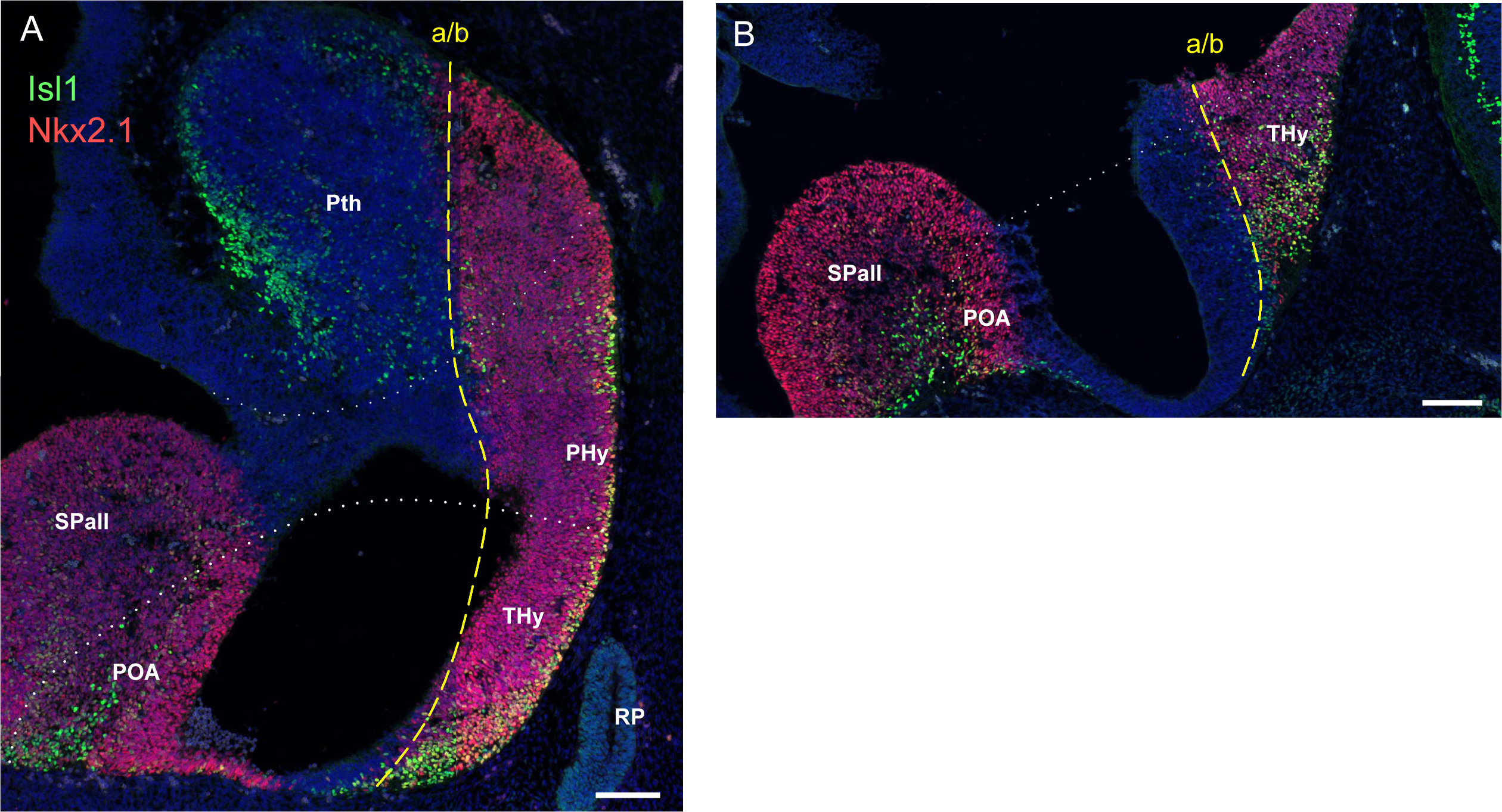
Double immunofluorescence of Isl1 (green) and Nkx2.1 (red) in sagital slices of a E10.5 mouse embryo. Nuclei are stained with DAPI. Panels go from lateral (A) to medial (B). Dotted white lines indicate the limits of anatomical regions. The yellow dotted line indicates the alar/basal limit (a/b).

Neurogenesis is well underway at E12.5, and at this stage the expression domain of Isl1 in specific forebrain subdivisions becomes clearer (Fig 3). Within the basal hypothalamus, Isl1 is expressed in the anterobasal area (ABas) and in more posterior regions of the tuberal hypothalamus, but not in the mamillary area (Mam). In the alar hypothalamus, Isl1 is found in the subparaventricular area (SPa) but seems to be absent from the paraventricular area (Pa). The Pth expresses Isl1 abundantly, both in the reticular nucleus (Rt) and Zona Incerta (ZI) as well as the POA and SPall (Fig. 3).

**Fig 3:**
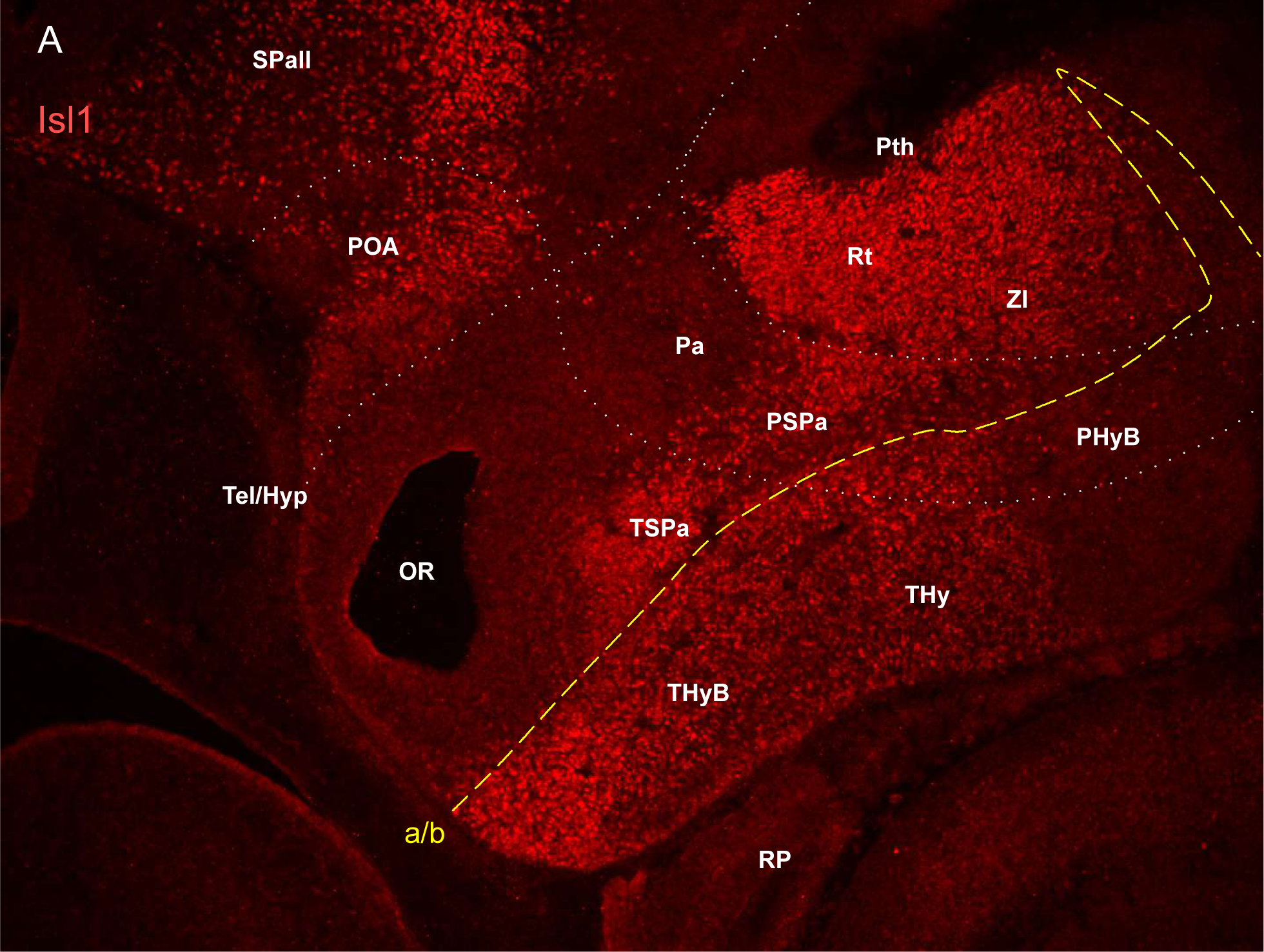
Immunofluorescence of Isl1 (red) in a sagital slice of a E12.5 mouse embryo. Dotted white lines indicate the limits of anatomical regions. The yellow dotted line indicates the alar/basal limit (a/b).

At E14.5, when neurogenesis is coming to a close, a comparison between the expression of Isl1 and Nkx2.1 shows that Isl1 is absent from the mamillary area, where Nkx2.1 is abundantly expressed (Fig 4). Similarly, the developing ventromedial hypothalamus (VMH) in the tuberal hypothamus is mostly devoid of Isl1+ cells but expresses Nkx2.1. A comparison with the expression of the POU-domain transcription factor Brn2 (POU3F2), which is a marker of the Pa, shows that Isl1 is completely absent from this area (Fig. 5). The HD transcription factor Pax6, which is not expressed in the hypothalamus, colocalises to a great extent with Isl1 in the Pth at E14.5 (Fig 6), something that was observed previously in E10.5 embryos (Mastick & Andrews, 2001). Lhx1 is also coexpressed with Isl1 in the Pth. In the hypothalamus, however, Lhx1 and Isl1 are expressed in mutally exclusive domains, with a sharp boundary between the Lhx1+ suprachiasmatic nucleus (SCN) and the Isl1+ ABas (Fig 7).

**Fig 4:**
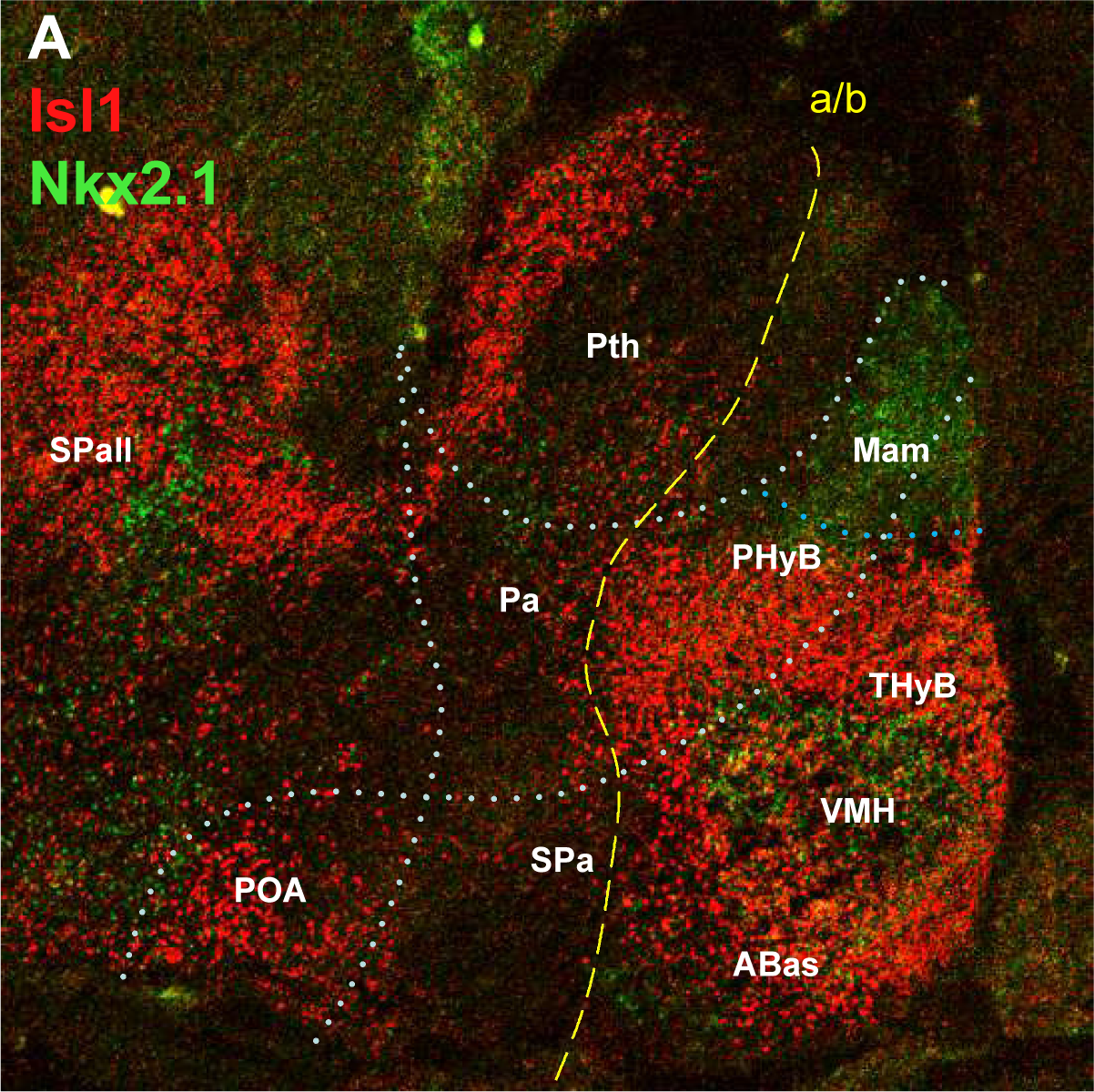
Double immunofluorescence of Isl1 (red) and Nkx2.1 (green) in a sagital slice of a E14.5 mouse embryo. Dotted white lines indicate the limits of anatomical regions. The yellow dotted line indicates the alar/basal limit (a/b).

**Fig 5:**
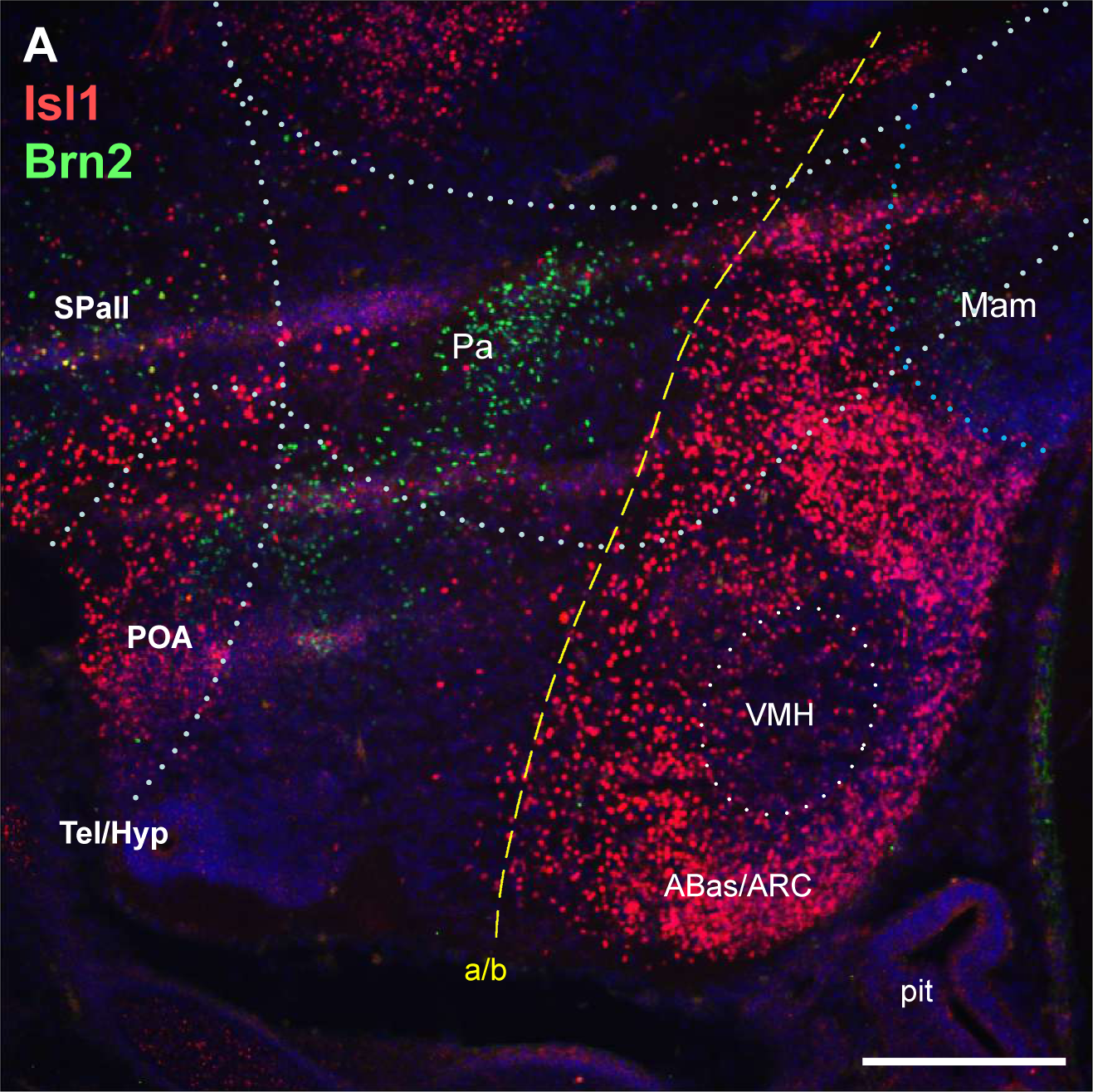
Double immunofluorescence of Isl1 (red) and Brn2 (green) in a sagital slice of a E14.5 mouse embryo. Dotted white lines indicate the limits of anatomical regions. The yellow dotted line indicates the alar/basal limit (a/b).

**Fig 6:**
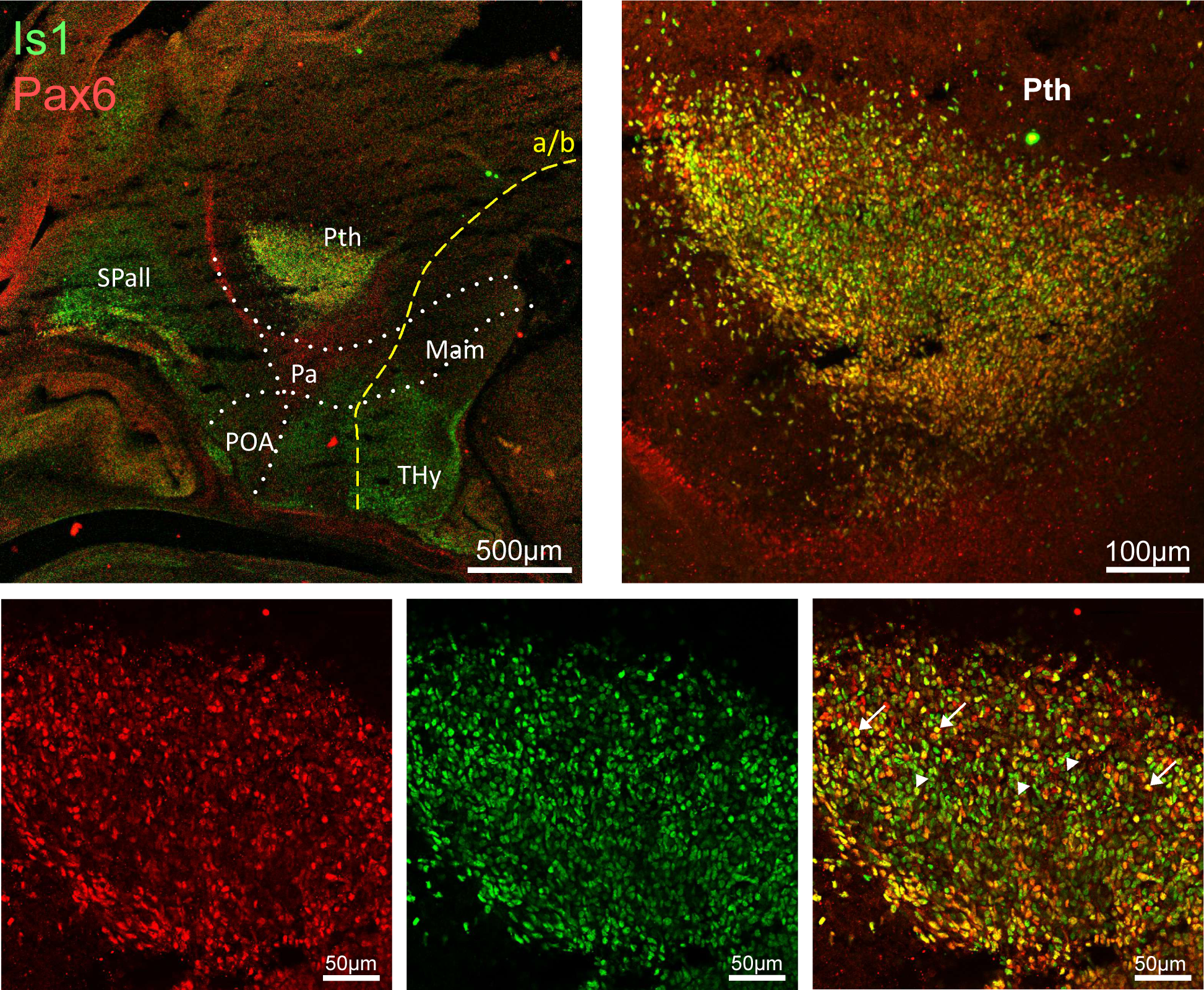
Double immunofluorescence of Isl1 (green) and Pax6 (red) in a sagital slice of a E14.5 mouse embryo. Insets show closeups of the prethalamic area. Dotted white lines indicate the limits of anatomical regions. The yellow dotted line indicates the alar/basal limit (a/b).

**Fig 7:**
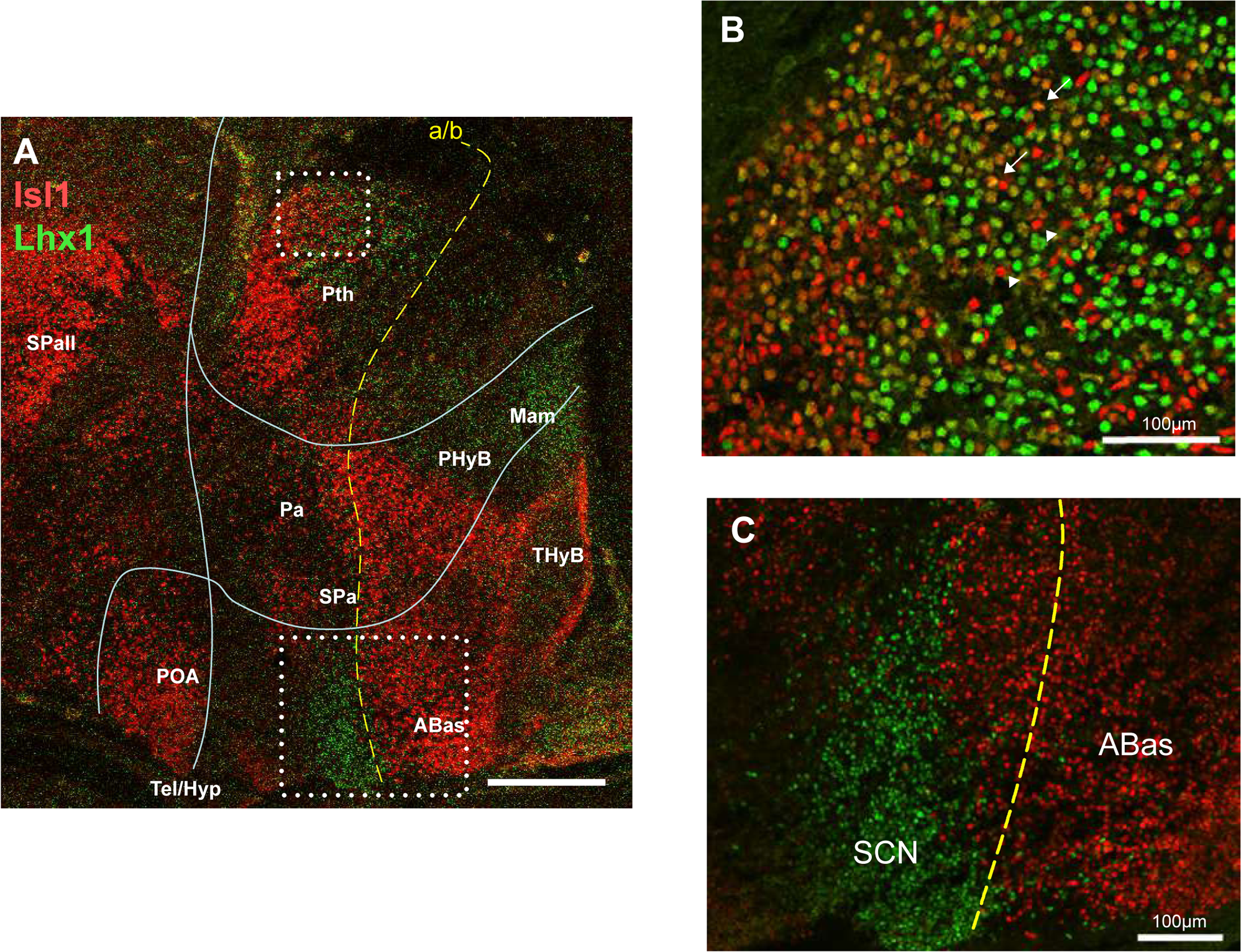
Double immunofluorescence of Isl1 (red) and Lhx1 (green) in a sagital slice of a E14.5 mouse embryo. Insets show closeups of the prethalamic and the SCN/ABas areas. Dotted white lines indicate the limits of anatomical regions. The yellow dotted line indicates the alar/basal limit (a/b).

### Colocalisation of Isl1 and TH in the forebrain

While mapping the distribution of Isl1+ cells in the forebrain, we noticed that the expression domain of this factor encompasses many of the areas where the dopaminergic molecular marker, tyrosine hydroxylase (TH), is known to be expressed (see for instance Marín et al, 2005). Indeed, at E10.5, TH immunofluorencence can be detected in the SPall and POA in the telencephalon, the Pth in the diencephalon and in part of the alar and basal hypothalamus, and most, if not all, of the TH+ neurons in these areas exhibit Isl1-positive nuclei (Fig 8). Only a small patch of TH+ cells, located in the Pa just anterior to the Pth, does not express Isl1 (Fig. 8).

**Fig 8:**
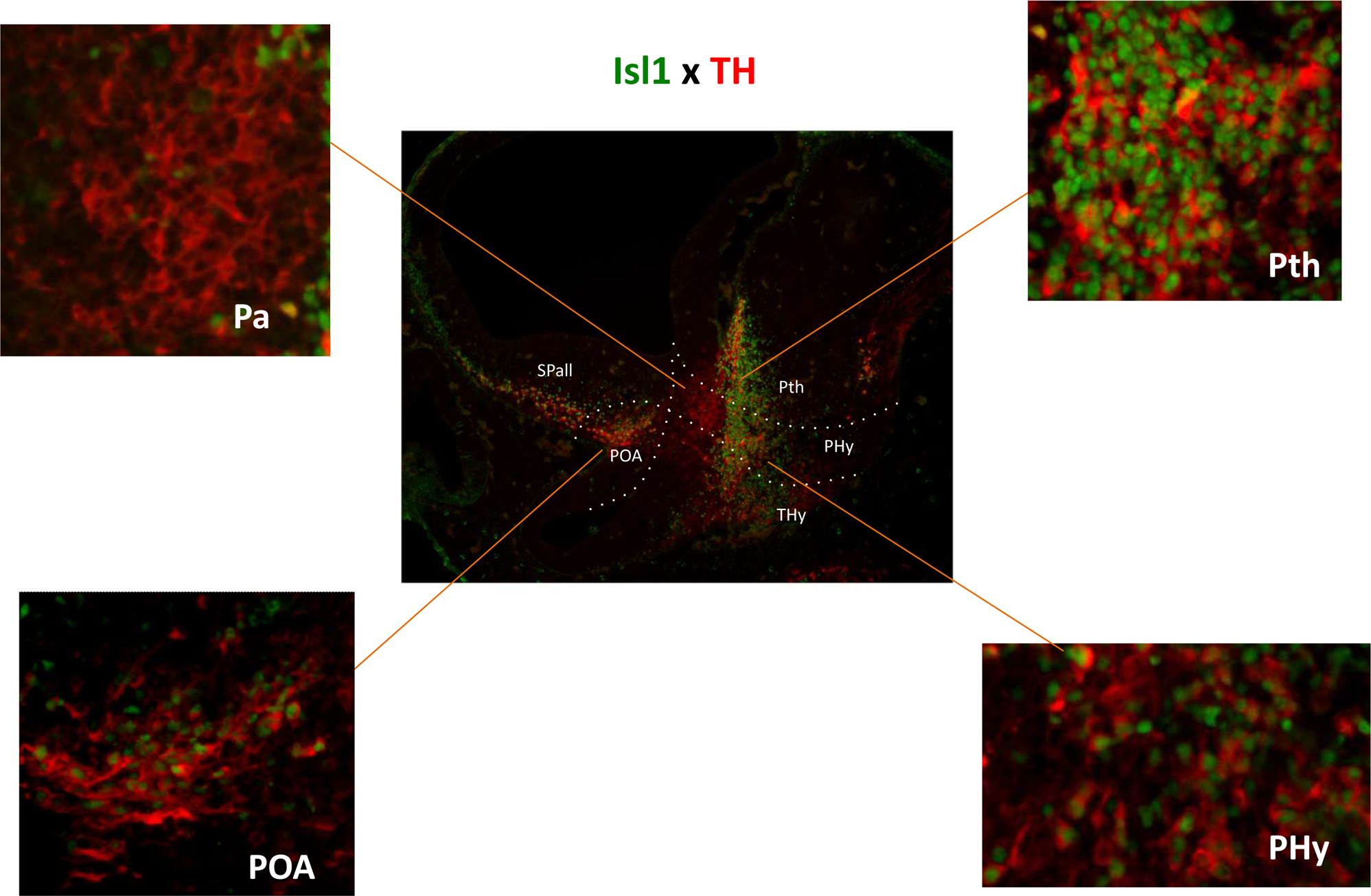
Double immunofluorescence of Isl1 (green) and TH (red) in a sagital slice of a E10.5 mouse embryo. Insets show closeups of the areas indicated by straight lines. Dotted white lines indicate the limits of anatomical regions. The yellow dotted line indicates the alar/basal limit (a/b).

To estimate the degree of colocalisation of Isl1 and TH during normal mouse development, we carried out double Isl1/TH immunofluorencence and scored the percentage of TH+ cells that where also Isl1+ in the POA, Pth and hypothalamic areas by confocal microscopy. The experiments were performed at early (E11.5) and late (E18.5) development, as well as adulthood.

At E11.5, the domain of expression of TH is somewhat continuous between the telencephalon, hypothalamus and diencephalon, and the typical dopaminergic neuronal groups are not easy to distinguish. We divided the TH+ domains in four groups, the POA (corresponding the future A15 group), the Pth (A13 group) and two areas in the hypothalamus, AH1 (encompassing the A14 and A12 in the AHA and ABas) and AH2, in the Pa. Figure 9 shows that the degree of colocalisation of TH and Isl1 is quite high in the POA, Pth and AH1, reaching nearly 90% in all these areas. The exception is AH2 in the Pa, where the a negligible percentage of TH+ cells are Isl1+ (Fig 9A,C).

**Fig 9:**
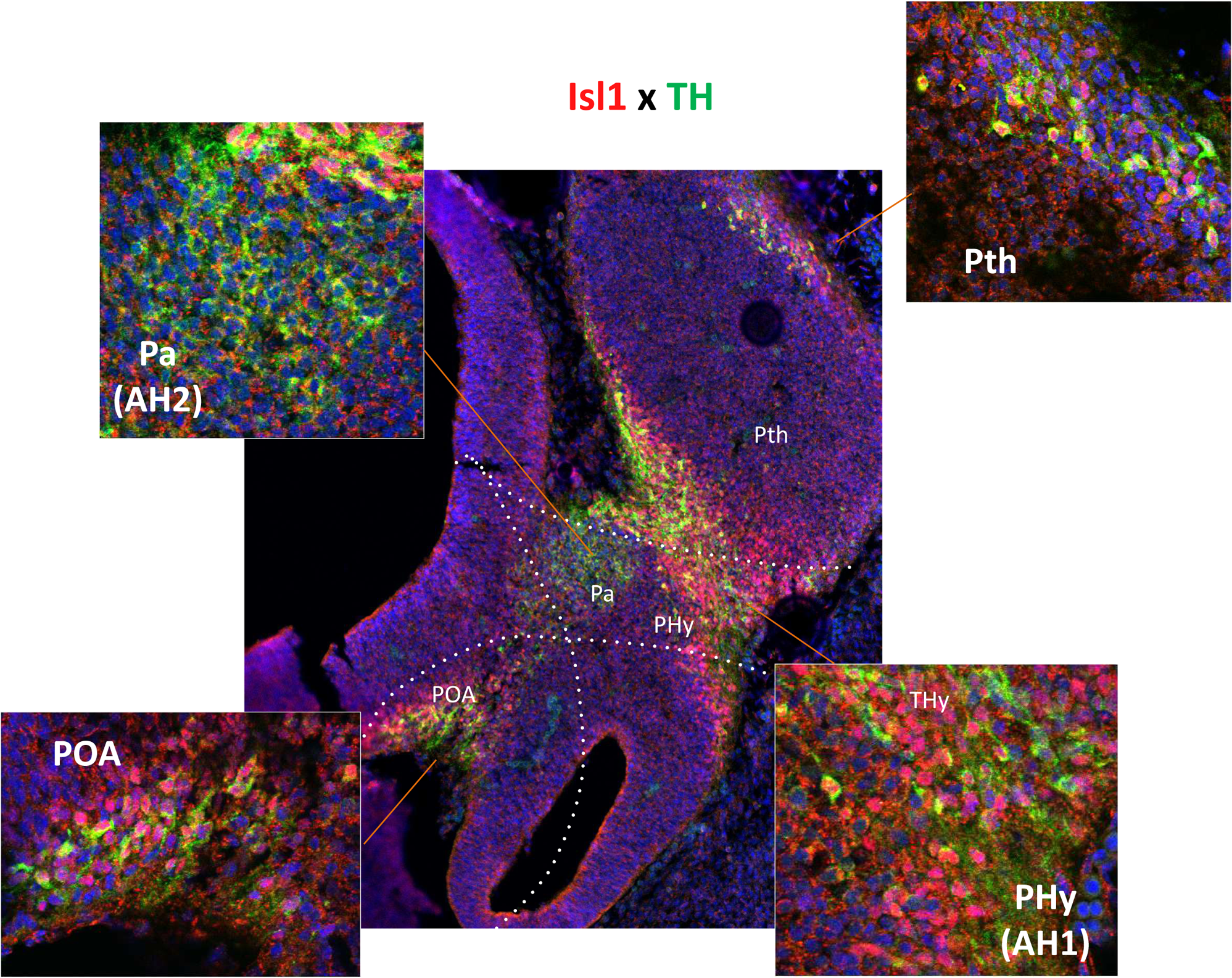
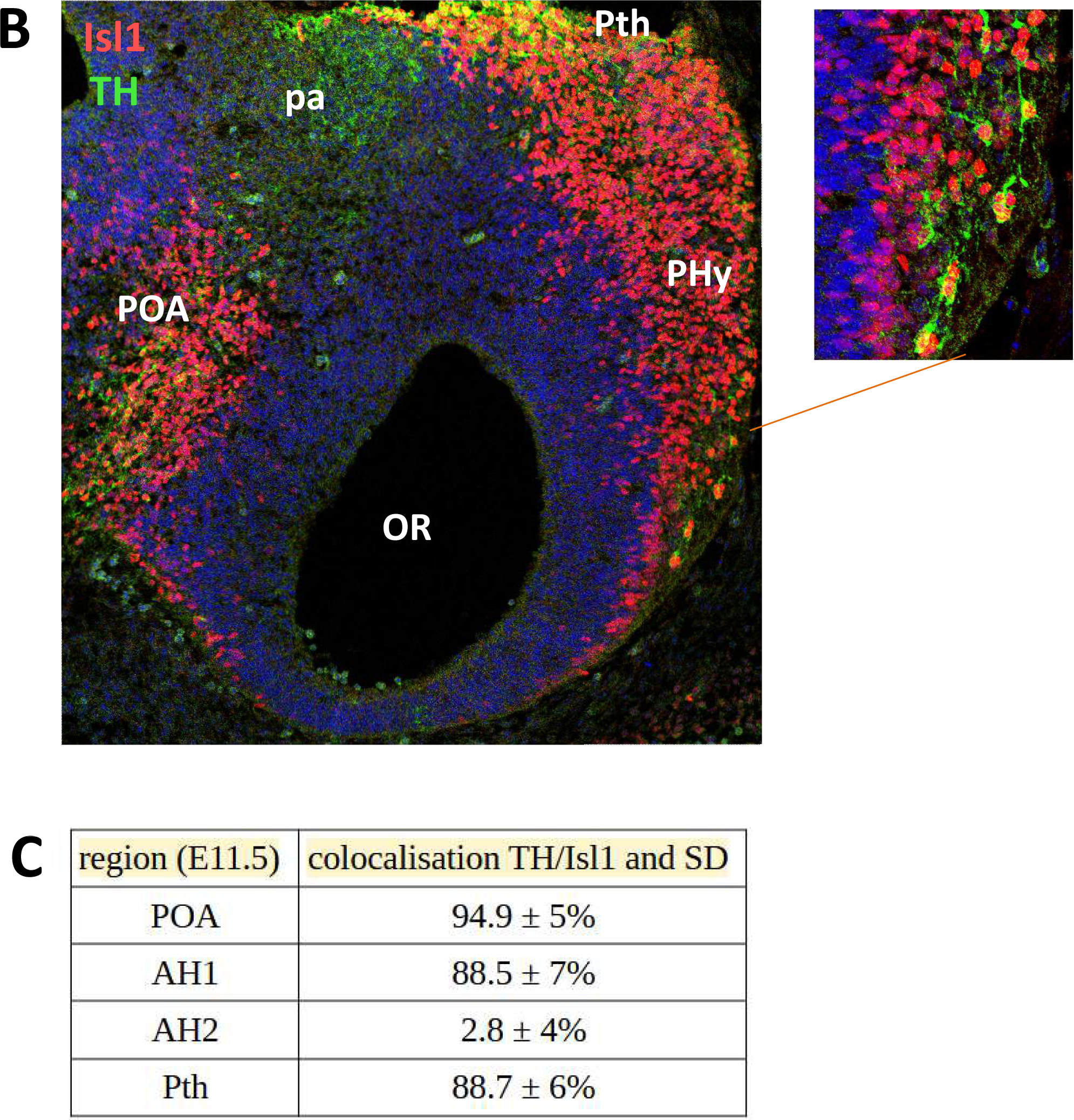
Colocalisation of Isl1 and TH at E11.5. Panels (A) and (B) show double immunofluorescence of Isl1 (red) and TH (green) in sagital slices of a E11.5 mouse embryo. Insets show closeups of the areas indicated by straight lines. Dotted white lines indicate the limits of anatomical regions. The yellow dotted line indicates the alar/basal limit (a/b). Panel (C) shows the quantifications of the percent of TH+ neurons with Isl1+ nuclei in each area with standard deviations.

In sagital sections of E18.5 embryos, the known dopaminergic groups, A15, A13 and A12, are more readly identified (Fig 10A). Also the A16 group, located in the olfactory bulb, and A11 in the posterior diencephalon can be identified. The A14 group, which is more scattered and located near the third ventricle of the hypothalamus, is best visualised in coronal sections, and we used these slices to perform the colocalisation studies (Fig 10B). The A14 group was divided into a rostral periventricular (A14-PeVR) and a mediocaudal (A14-PeMC) subgroup. The colocalisation results indicate that TH neurons belonging to the ARC/A12, Pth/A13, POA/A15 have high levels of Isl1 coexpression, specially the A13 group, with over 80% TH/Isl1 neurons (Fig 10B-C). The A14-PeVR displayed around 70% of TH/Isl1 colocalisation, while the A14-PeVMC was much lower at around 11%, similar to the Pa group. The A11 and A16 dopaminergic groups, in contrast, do not express Isl1 at all (Fig 11 and data not shown).

**Fig 10:**
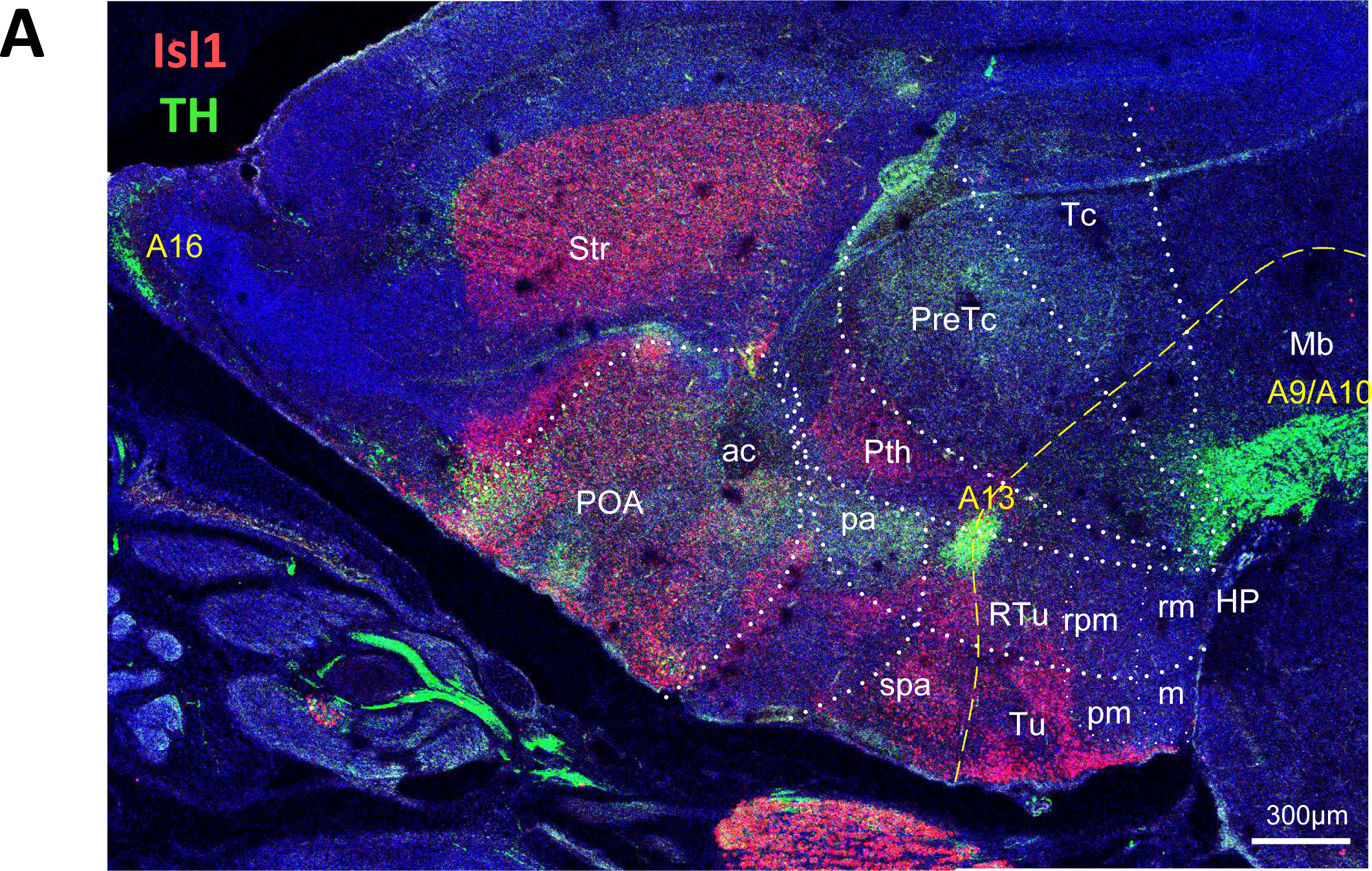
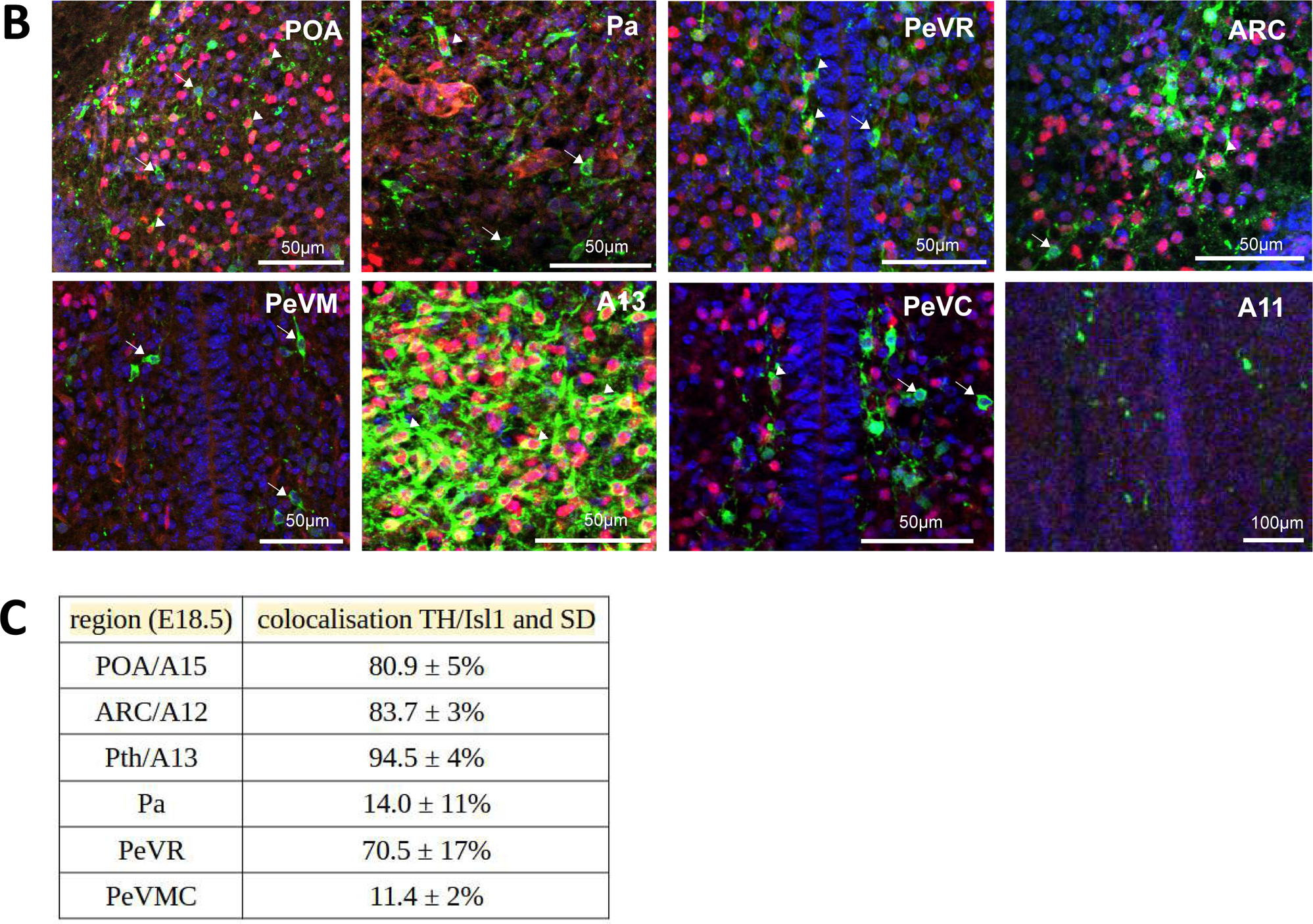
Colocalisation of Isl1 and TH at E18.5. Panel (**A**) shows double immunofluorescence of Isl1 (red) and TH (green) in a sagital slice of a E18.5 mouse brain. Dotted white lines indicate the limits of anatomical regions. The yellow dotted line indicates the alar/basal limit (a/b). Panel (**B**) shows coronal images of the areas used for colocalisation estimations. Arrowheads indicate TH+/Isl1+ neurons while arrows point to TH+ cells that do not express Isl1. Panel (**C**) shows the quantifications of the percent of TH+ neurons with Isl1+ nuclei in each area with standard deviations.

**Fig 11:**
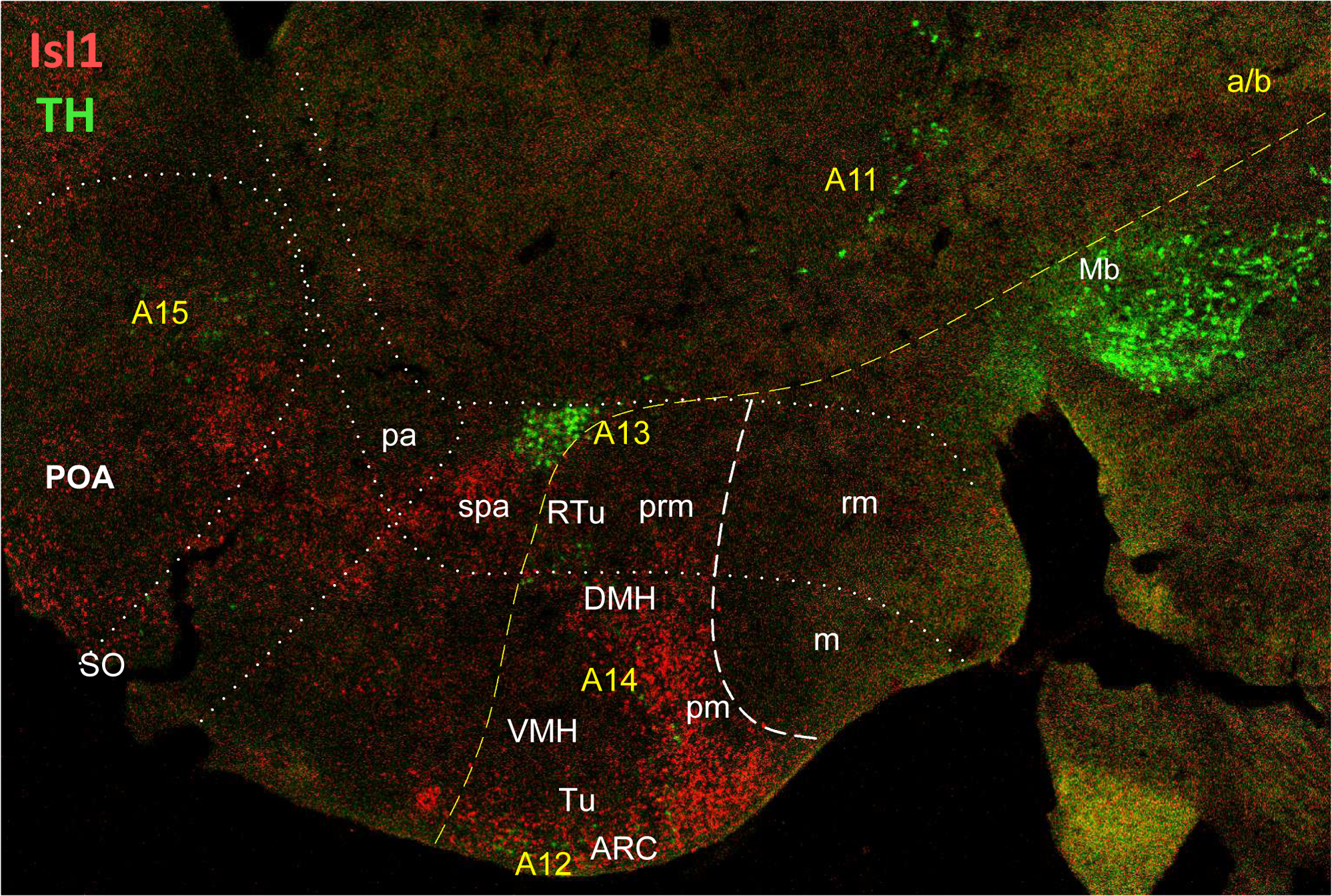
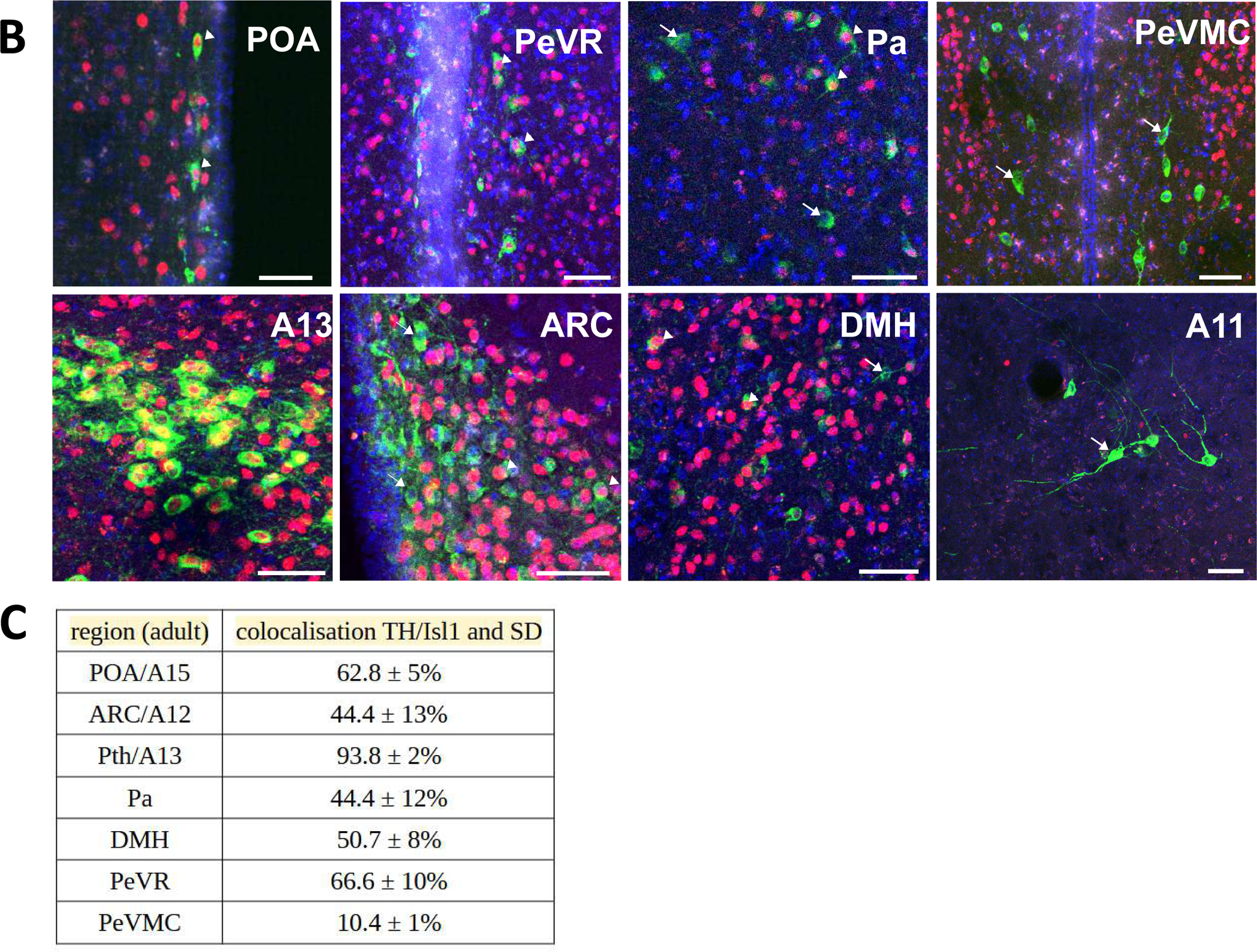
Colocalisation of Isl1 and TH at adulthood. Panel (**A**) shows double immunofluorescence of Isl1 (red) and TH (green) in a sagital slice of the brain of a two-month old adult mouse. Dotted white lines indicate the limits of anatomical regions. The yellow dotted line indicates the alar/basal limit (a/b). Panel (**B**) shows coronal images of the areas used for colocalisation estimations. Arrowheads indicate TH+/Isl1+ neurons while arrows point to TH+ cells that do not express Isl1. Panel (**C**) shows the quantifications of the percent of TH+ neurons with Isl1+ nuclei in each area with standard deviations.

At adulthood, sagital sections again reveal the classical dopaminergic groups (Fig 11A). In addition, dopaminergic neurons are found in the Pa and in the dorsomedial hypothalamus (DMH), a population that might also be considered as part of the A14 group. Double immunofluorencence experiments performed in coronal sections show that, at this stage, only the Pth/A13 population of TH neurons coexpresses Isl1 at very high levels, reaching over 90% of colocalisation, followed by the POA/A15 and the A14-PeVR, with around 60% (Fig 11B-C). The ARC/A12, DMH and Pa groups display over 40% colocalisation, while the A14-VMC group has a very low level of TH/Isl1 coexpression at around 10%. The A11 group does not include any Isl1+ neurons. These results are summarised in Figure 12.

**Fig 12:**
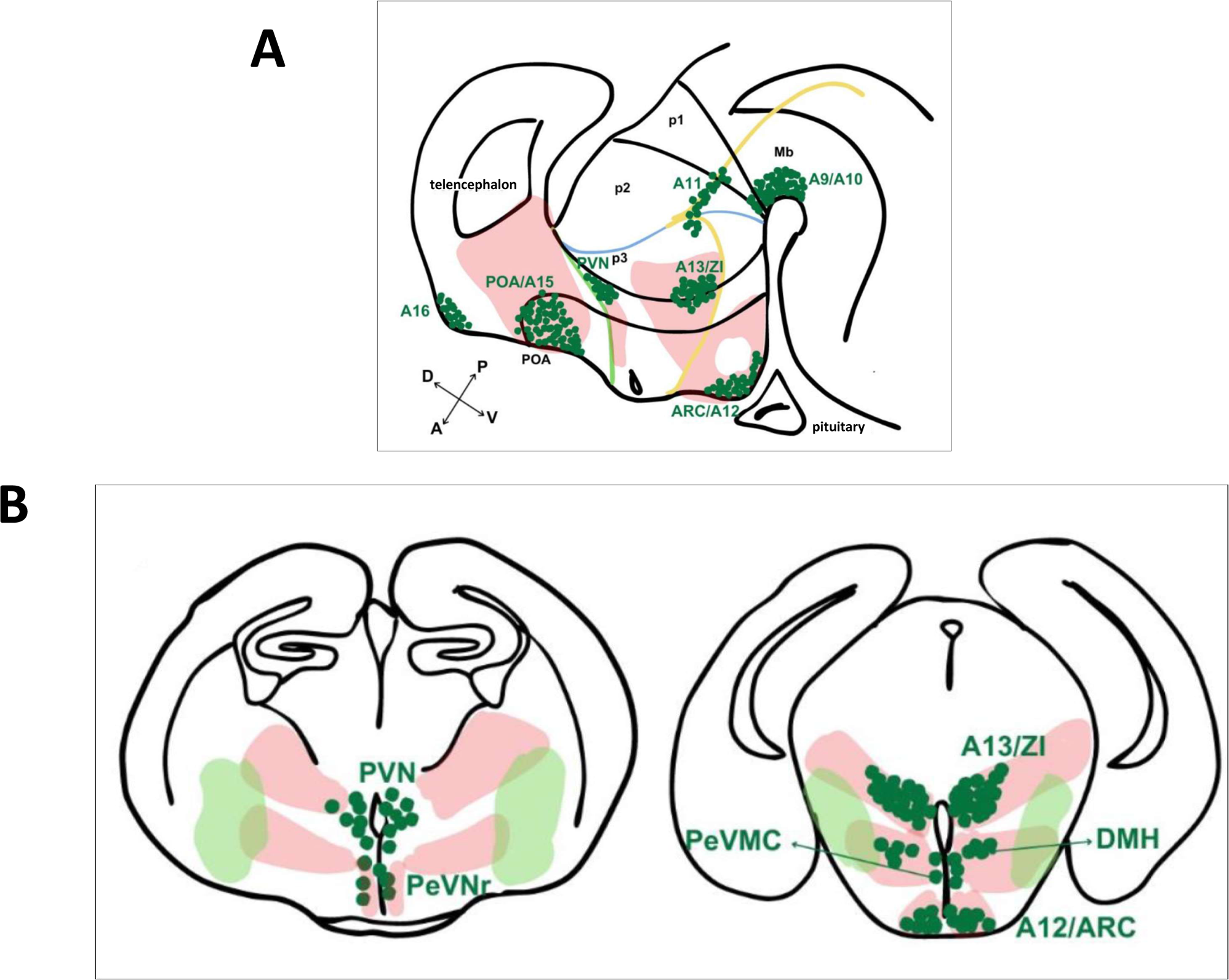
Schematics of the pattern of expression of Isl1 and TH in a sagital slice of a E14.5 embryo (**A**) and in transversal slices of an adult mouse brain (**B**). Note that most TH neuronal groups of the forebrain lie within the territory of Isl1 expression.

## Discussion

The expression of Isl1 has been studied by immunofluorescence in the forebrain of other tetrapods, including the frog, *Xenopus laevis* (Moreno et al, 2008; Domínguez et al, 2013, 2014, 2015), the turtle, *Pseudemys scripta* (Moreno et al, 2012) and a urodele amphibian (*Pleurodeles waltl*) together with two lungfishes, *Neoceratodus forsteri* and *Protopterus dolloi* (Moreno et al, 2018). In all of these animals, the expression of Isl1 is very similar to the one in the mouse. Thus, in all species analysed, Isl1+ neurons populate the following regions: i) the subpallium and the preoptic region in the telencephalon; ii) the suprachiasmatic area in the alar hypothalamus, which seems to be the anatomical equivalent of the subparaventricular area in the mouse; iii) most of the tuberal hypothalamus and iv) the prethalamus. On the other hand, Isl1 is absent from the paraventricular area, in the alar hypothalamus, and from the mamillary area in the basal hypothalamus in all tetrapods analysed. An important detail in the mouse is the total absence of Isl1 in the suprachiasmatic nucleus (SCN), which is delineated by Lhx1 expression in the alar hypothalamus. The SCN is, however, part of the paraventricular area in the mouse, where Isl1 is absent in all species. In addition to work done in tetrapods, mapping of Isl1 expression in the zebrafish and non-teleost actinopterygian fishes reveal a similar pattern of Isl1 distribution in the forebrain (Schredelseker and Driever, 2020; López et al, 2021; Lozano et al, 2023). Thus, Isl1 expression is conserved in vertebrates and characterises the anteriormost part of the forebrain, from the ventral telencephalon and hypothalamus to the diencephalic prosomere 3. Isl1 expression in more posterior levels of the brain is mostly restricted to motor nuclei in the mid and hindbrain.

A pioneering work by Grimm et al (2004) compared the transcriptome of dopaminergic cell populations obtained by laser-capture and identified *Isl1* mRNA in the A13 group but not in midbrain dopaminergic groups. More recent single-cell RNA sequencing analyses of the mouse hypothalamus have also identified *Isl1* as a molecular marker of dopaminergic neurons in the forebrain, in particular in the prethalamus (Hook et al, 2018; Romanov et al., 2020; Kim et al, 2020). Using a Isl1-Cre transgene and the Ai14 lineage tracer, Romanov at al (2020) have mapped the contribution of Isl1 expressing neurons to the dopaminergic lineage and found that, at E14.5, many TH+ neurons of the ARC/A12, Pth/A13, POA/A15, A14-PeVR and DMH expressed the tomato-td2 reporter induced by Isl1-Cre (Extended Data Fig. 9 in Romanov et al, 2020). In their experiments, the colocalisation of tomato-td2 and TH is not complete, not even in the Pth/A13, where we observed the highest degree of colocalisation, which may reflect the limitations of the genetic lineage tracer methodology. Interestingly, Lee et al (2016) used to immunofluorescence to estimate Isl1/TH coexpression at around 70% at E16.5 and 25% at P56, which is consistent with our observation of a general decline in the percentage of Isl1/TH colocalisation in most areas. Altogether, our study provides spatial and temporal information as to the degree of Isl1 and TH colocalisation in the mouse forebrain. Thus, at the beginning of neurogenesis, Isl1 is found in all dopaminergic neurons in the basal telencephalon, hypothalamus and prethalamic regions, except for a small population of TH+ cells located in the paraventricular area of the alar hypothalamus. In later development and adulthood, Isl1 is still expressed in a majority of dopaminergic neurons of the Pth/A13 group (90%) and in a moderate percentage (40-50%) in the ARC/A12, POA/A15 and areas associated with the A14 group, like the rostral part of the periventricular region (A14-PeVR) and the DMH (Fig. 12).

Little is known about the transcription factors that control dopaminergic cell fate in the murine forebrain. The bHLH factor Ascl1 is necessary for the differentiation of A12, A13, A14 and A15 groups (Romanov et al, 2020). Specifically for the A13 group, transcription factors Dlx1/2 (Andrews et al, 2003) and Arx (Sunnen et al, 2014) are also necessary for TH expression. Mice mutant for the transcription factors Onecut 1 (Hnf6) and Onecut 2 have reduced numbers of TH neurons of the A13 group (Espana et al, 2012), and Orthopedia (Otp) is needed in the A11 group (Ryu et al, 2007). Using a mouse conditional knockout approach, Lee et al (2016) showed that *Isl1* is not necessary to sustain TH expression in the ARC/A12 group. In the zebrafish, *isl1* is expressed in prethalamic TH neurons, and the injection of embryos with morpholinos targetting *isl1* caused the disappearance of TH neurons from the prethalamus, while TH expression in the hypothalamus remained unaltered (Filippi et al, 2014). It remains to be seen whether Isl1 is important for dopaminergic neuron differentiation in the mouse forebrain.

In summary, in this work we describe Isl1 expression in the mouse forebrain and dopaminergic neurons. We hope it may provide a framework for the study of the fuction of this transcription factor in the mammalian brain.

## Abbreviations

a/b: alar/basal limit
ABas: anterobasal area
Arc: arcuate nucleus
DMH: dorsomedial hypothalamus
Mam: mamillary area
Mb: midbrain
Pa: paraventricular area
PeVR: rostral periventricular area
PeVMC: ventrocaudal periventricular area
PHy: peduncular hypothalamus
PHyB: basal plate of the peduncular hypothalamus
POA: preoptic area
PVN: paraventricular nucleus
p1: prosomere 1
p2: prosomere 2
p3: prosomere 3
Pit: pituitary glando
rm: retromamillary area
RP: Rathke’s pouch
Rt: reticular nucleus
Pth: prethalamus
RTu: retrotuberal area
SCN: suprachiasmatic nucleus
SO: supraoptic nucleus
SPa: subparaventricular area
SPall: subpallium
Str: striatum
Tel: telencephalon
THy: terminal hypothalamus
THyB: basal plate of the terminal hypothalamus
Tu: tuberal area of hypothalamus
VMH: ventromedial hypothalamus
ZLI: *Zona Limitans Intrathalamica*
ZI: *Zona Incerta*

## Author constributions

ACC designed and performed most experiments; MR helped designed the study and provided materials; FSJS designed the study, performed some experiments and wrote the manuscript with input from the other authors. All authors approved the final manuscript.

## Acknowledgements

This work was supported by grants from the Agencia Nacional de Promoción Científica y Tecnológica (ANPCyT, BID PICT 2018-03947 and PICT 2020-SERIEA-03451). We would like to thank Lucía Franchini and Estela Muñoz for antibody gifts. ACC was supported by PhD fellowships by CONICET and ANPCyT, Argentina. MR and FSJS are CONICET career investigators, Argentina.

